# Biomolecular condensates can both accelerate and suppress aggregation of α-synuclein

**DOI:** 10.1101/2022.04.22.489149

**Authors:** Wojciech P. Lipiński, Brent S. Visser, Irina Robu, Mohammad A. A. Fakhree, Saskia Lindhoud, Mireille M. A. E. Claessens, Evan Spruijt

## Abstract

Biomolecular condensates present in cells can fundamentally affect the aggregation of amyloidogenic proteins and play a role in the regulation of this process. While liquid-liquid phase separation of amyloidogenic proteins by themselves can act as an alternative nucleation pathway, interaction of partly disordered aggregation-prone proteins with pre-existing condensates that act as localization centers could be a far more general mechanism of altering their aggregation behavior. Here, we show that so-called host biomolecular condensates can both accelerate and slow down amyloid formation. We study the amyloidogenic protein α-synuclein and two truncated α-synuclein variants in the presence of three types of condensates composed of non-aggregating peptides, RNA or ATP. Our results demonstrate that condensates can dramatically speed up amyloid formation when proteins localize to their interface. However, condensates can also significantly suppress aggregation by sequestering and stabilizing amyloidogenic proteins, thereby providing living cells with a possible protection mechanism against amyloid formation.

## Introduction

With increasing life expectancy, neurodegenerative diseases involving pathological amyloid formation are becoming alarmingly common. Misfolded and aggregated proteins may accumulate during the lifetime of a cell when these are not refolded or cleared by protein quality control machinery (1). Such accumulation can hamper regular cell operations, eventually leading to cell death and, at the organ level, to various neurodegenerative diseases including Alzheimer’s and Parkinson’s disease (2–4). For many years researchers have studied the origins, mechanism, and toxicity of protein aggregation, in order to develop effective therapies, but many aspects of the mechanism of protein aggregation remain incompletely understood (5–8). The quest for new therapies is impeded by the fact that processes inside cells take place in a complex environment that is difficult to reproduce in vitro (9). By contrast, the vast majority of protein aggregation studies are carried out with synthetic peptides or proteins fragments in dilute, homogeneous and well-mixed solutions (10, 11).

One of the most striking features that distinguishes the complex intracellular environment from the protein solutions often used in vitro is the presence of condensates formed by liquid-liquid phase separation (LLPS) of biomolecules into so-called membraneless organelles (MLOs) (12–15). These compartments are usually liquid-like, highly concentrated droplets of proteins and nucleic acids. Examples of such organelles include nucleoli (16) and Cajal bodies (17) in the nucleus, and stress granules in the cytoplasm (18). The main difference between MLOs and membrane-bound compartments is the lack of a physical barrier between the organelle and the surrounding solution. This results in the ability to exchange components with the environment, to undergo fusion and to respond to environmental changes by rapid formation/dissolution (19).

Knowing that MLOs contain proteins at very high concentrations and that proteins that undergo LLPS and proteins that partition into liquid droplets often feature low-complexity domains (13, 20), a characteristic that is also common for amyloidogenic proteins, it becomes evident that the presence of biological condensates could drastically affect the aggregation process. For various phase-separating proteins it has been suggested that prior condensation into liquid droplets can promote conformational changes within the disordered region leading to the formation of gel-like structures or amyloid-like aggregates. Such a process has been observed, for instance, for hnRNPA1 (18), FUS (21) or Tau (22, 23). Recently, it has been shown that also α-synuclein (αSyn), one of the archetypical amyloid-forming proteins, can undergo LLPS under PEG-based crowded conditions, and that the condensed α-synuclein droplets may facilitate aggregation (24–26). However, it remains unclear if LLPS of α-synuclein LLPS is also likely to happen in living cells, as α-synuclein is known to interact with many components inside the cell, including membranes, the cytoskeleton and other proteins (27, 28), which may suppress the concentration of free α-synuclein and prevent the formation of homotypic α-synuclein condensates.

Nonetheless, there is also another, more general way by which LLPS can affect protein aggregation, which is also relevant for proteins that are present in cells at low concentrations. Condensates can concentrate guest biomolecules, including amyloidogenic proteins, by partitioning or interfacial adsorption, and provide a distinct chemical environment in which the stability and reactivity of biomolecules may be impacted. This can alter the kinetics of protein aggregation in multiple ways (29–31). An enhanced local concentration of amyloidogenic proteins may result in acceleration of the aggregation process, according to the law of mass action (32). However, one has to take into account that the local environment of the condensed liquid may promote protein conformations that do not undergo aggregation as readily as ones dominating in the surrounding solution. This has been observed for amyloid-β(1–42) (33) and may occur also for other amyloidogenic proteins. Finally, aggregating proteins could accumulate at the interface between condensate and the surrounding cytosol, potentially resulting in alternative, interfacial aggregation pathways, analogous to what has been observed for lipid vesicles and solid surfaces (34). In general, accumulation of amyloidogenic proteins at an interface can alter the kinetics of aggregation in two ways: an increased local concentration leads to faster aggregation, and an altered conformation of molecules bound to the interface can either stabilise free monomers or promote their transformation into fibrils.

Accumulation of αSyn at the interface has been reported to accelerate aggregation, e.g. for exosomes and SUVs (small unilamellar vesicles) (35). By contrast, aggregation was slowed down when SUV were present in large excess over αSyn and most protein monomers were trapped in a stable configuration at the surface of SUVs and there was no free monomeric αSyn in solution (36). Similar effects have been observed for SUVs and LUVs (large unilamellar vesicles) composed of mixtures of anionic, cationic and neutral lipids (37). The effect of lipid membranes on αSyn is largely dependent on the lipid/protein ratio, but also on the chemical structure of lipid, mutations in the protein chain and probably also on the size (and thus the curvature) of the vesicles or surface defects associated with curvature (38). Interestingly, it has been suggested that the presence of membranes can even induce fibril dissociation, by stabilising monomers and depleting the solution of free protein (39).

While there is ample evidence that biomolecular condensates can fundamentally alter protein distributions in vitro and in living cells by concentration, exclusion or interfacial localization, a systematic investigation of the effects that pre-existing condensates have on protein aggregation is lacking. Here, we study the consequences of “inert” model condensates (coacervates) on the aggregation of α-synuclein. Our model condensates are inert in the sense that they are not composed of, or dependent on, the aggregating protein, and they do not undergo any form of liquid-to-solid transition themselves. The goal of using these model condensates is to investigate if pre-existing biomolecular condensates can have a generic effect on protein aggregation, by means of concentration, exclusion or stabilization of disordered confirmations.

In the experiments, we use full-length α-synuclein and two truncated variants to better understand which protein domains are responsible for specific behaviours. Three different coacervates were investigated as model condensates, and their selection was guided by well-defined coacervate models reported in literature of RNP granules containing RNA and arginine-rich peptides (40–42), heterotypic condensates containing unstructured polypeptides (43), and active droplets containing small molecules (44, 45). We show that depending on the composition of the condensed phase, amyloidogenic protein can partition into the droplets, remain excluded or accumulate at the interface. We find that full-length αSyn accumulates and aggregates preferentially inside two of the complex coacervate droplets and accumulates and aggregates at the interface of another type. Accumulation of full-length αSyn either inside or at the interface of coacervates always leads to enhanced aggregation compared to a homogeneous solution. Truncated variants of αSyn were typically excluded from the coacervate droplets and aggregated at comparable rate or slower than in homogenous solution. Interestingly, the shortest variant, which only contains the β-sheet forming region and which normally has the fastest maximum rate of aggregation, is also accumulated inside two coacervates, but aggregates significantly slower than in solution. This demonstrates that sequestration of amyloidogenic proteins inside condensates can speed up aggregation by enhancing local concentrations in some condensates, but slow it down in others due to a stabilisation of the monomeric form of the protein.

## Results and Discussion

### Properties of selected α-synuclein variants

We selected three αSyn variants with different net charge and length of the intrinsically disordered region (fig. 1A): wild-type, full-length α-synuclein (FL-αSyn), a truncated variant depleted of the negatively charged, disordered C-terminal domain (αSyn-1-108) and a relatively hydrophobic short peptide from the non-amyloid-β component of the protein, which is the part that is responsible for β-sheet formation in aggregates (NACore peptide, αSyn-68-78). While having different physicochemical properties, these variants are all able to aggregate into amyloids (fig. 1B), and the kinetics of their aggregation can be described by a classical nucleation and growth model (46), including primary and secondary nucleation (fig. 1C).

**Fig. 1.**
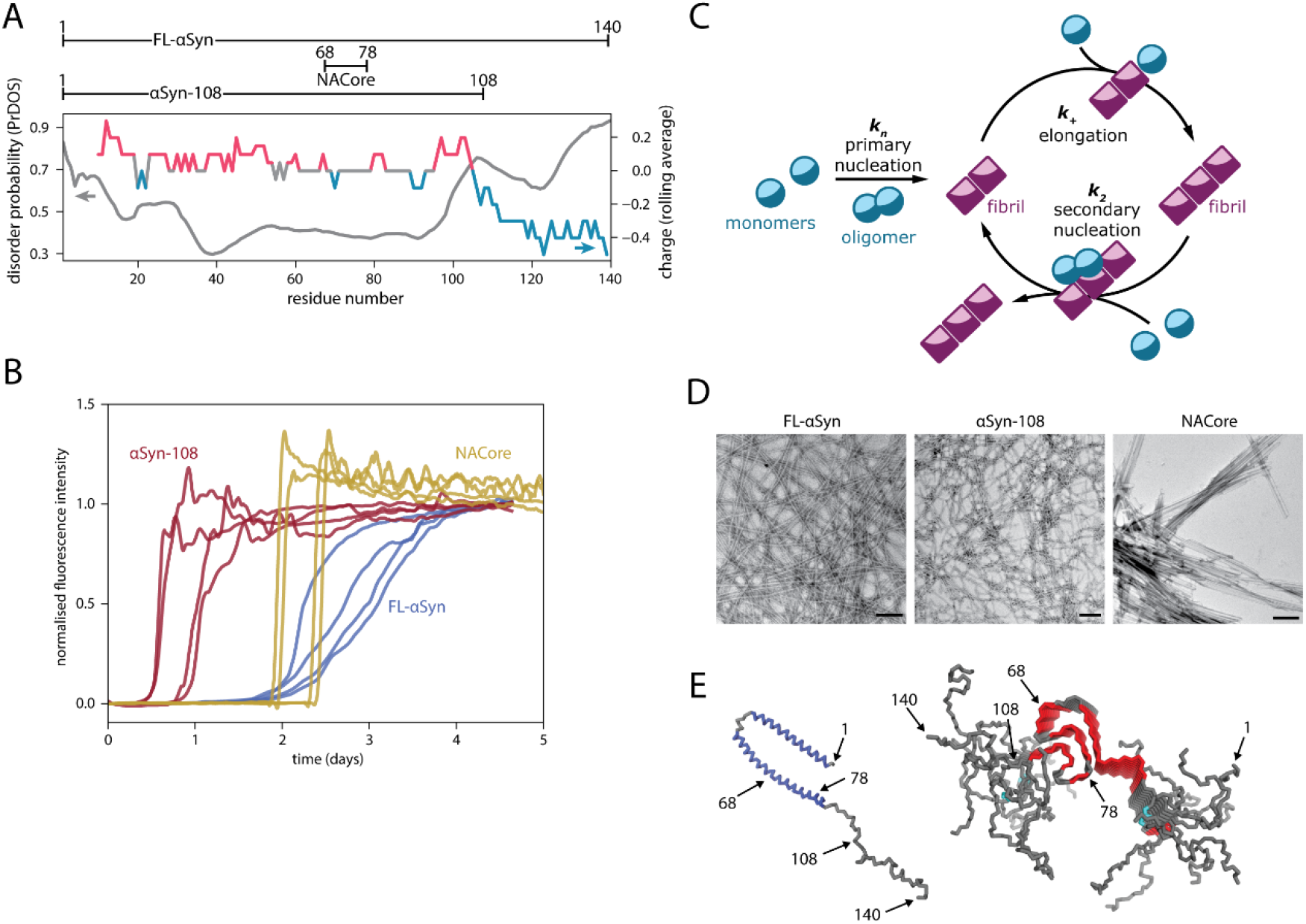
Amyloidogenic α-synuclein variants used in this study. (**A**) Variants of α-synuclein used in the study and predicted disorder along the protein chain (PrDOS, in grey) (*47*) and distribution of charged residues (in colour). For comparison of predicted disorder using different online tools please check supplementary information, fig. S7. (**B**) Aggregation traces (normalised thioflavin T fluorescence intensity) for various α-synuclein variants (recorded for 40 μM concentration of FL-αSyn and αSyn-108, and for 160 μM concentration of NACore). (**C**) Schematic depiction of the basic protein aggregation cycle model used in this study. (**D**) TEM images of fibrils formed by studied variants Scale bar = 200 nm. (**E**) FL-αSyn conformation when bound to lipids (left, PDB ID: 1XQ8) and stacked in amyloid fibrils (right, PDB ID: 2N0A), relevant residues are indicated.

All variants form fibrillar aggregates, as confirmed by TEM (fig. 1D). For each variant, we determined the concentration at which complete aggregation, defined as reaching plateau of ThT intensity, was reached in less than five days, and these concentrations were used in further experiments unless stated otherwise (40 μM for FL-αSyn and αSyn-108, and 160 μM for NACore).

### Partitioning of α-synuclein into coacervate droplets

We investigated the localisation of selected α-synuclein variants in 3 coacervate systems, which may serve as very basic models of biological condensates: (i) (RRASL)3 peptide/polyuridilic acid (RP3/polyU) (48, 49), (ii) poly-D,L-lysine/poly-D,L-glutamate (pLys/pGlu) (50, 51), and (iii) poly-L-lysine/ATP (pLys/ATP) (44, 45) (fig. 2). All these systems phase separate upon mixing (poly)cationic with a (poly)anionic components and form micrometre-size droplets that fuse into larger droplets over time, but remain liquid over the course of several days. In addition, all droplets have been shown to take up or exclude a wide range of biomolecules and complexes (40, 52–54), and are thus expected to influence the aggregation of α-synuclein. RP3/polyU coacervates have been suggested as a model for RNP granules, which are typically compose of RNA and arginine-rich peptides and disordered proteins. It is interesting to note that it contains a relatively short cationic component (RP3 peptide) and a long anionic component (polyU RNA), and is thus expected to interact weakly with negatively charged guest molecules. pLys/ATP coacervates also have peptide-nucleotide composition, and has been used as active droplet mimic. Unlike RP3/polyU coacervates, pLys/ATP coacervates contain a long cationic and a short anionic component, typically resulting in a positive surface potential when prepared at equal charge, and a strong interaction with negatively charged guest molecules. Finally, pLys/pGlu coacervates are composed of large unstructured cationic and anionic peptides, and have been used as model for IDP-based condensates without significant secondary structure formation upon condensation. Three different systems were selected as we expected different interactions between α-synuclein variants and different coacervate systems, which we hope may yield generalizable, physicochemical insight into the influence of the coacervate phase on protein aggregation.

**Fig. 2.**
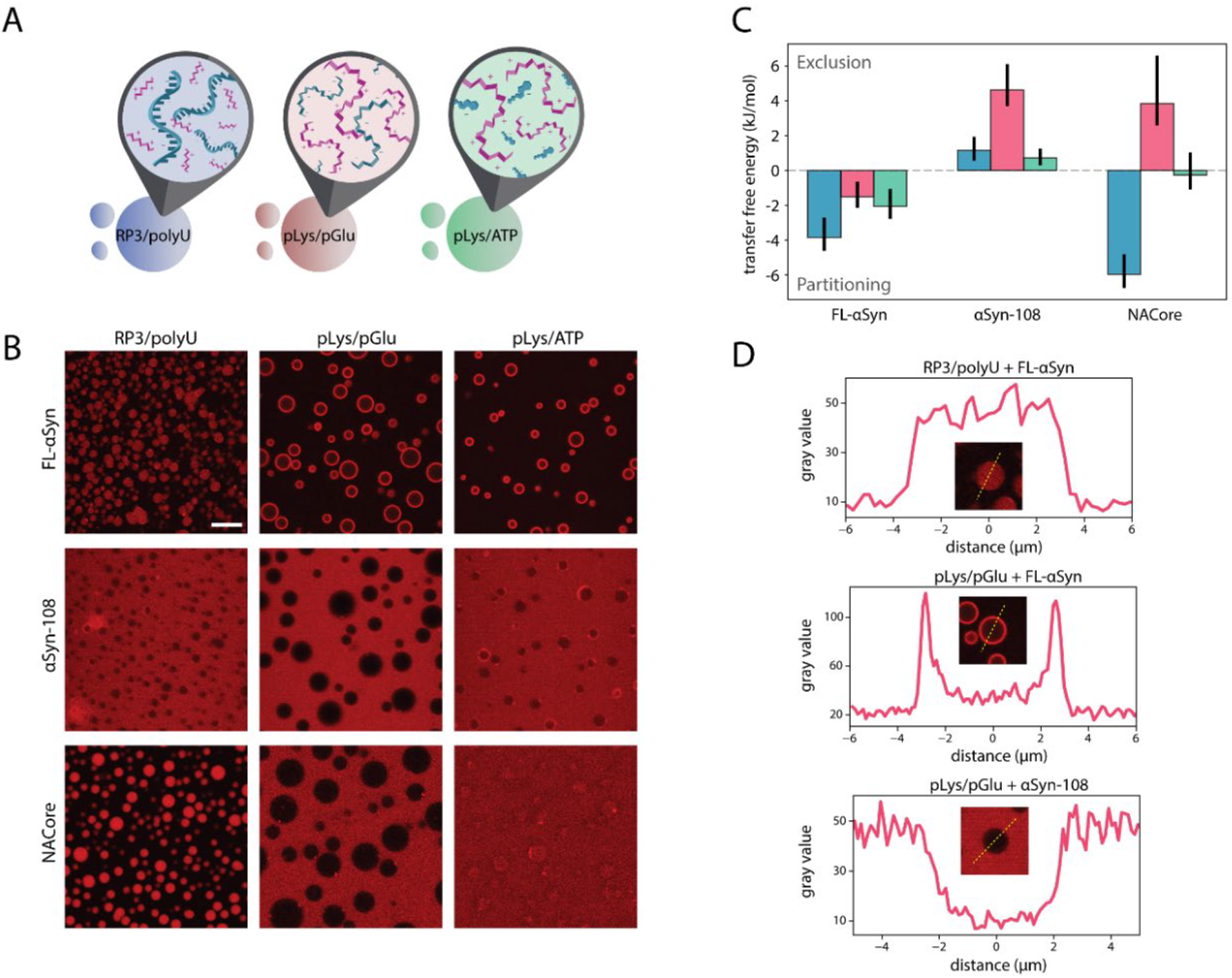
Coacervate systems and interactions with α-synuclein variants. (**A**) Schematic depiction of coacervate systems used in the study. (**B**) Confocal microscope images of coacervate systems with labelled α-synuclein variants, colorised artificially (Alexa Fluor 647-labelled S9C-FL-αSyn and S9C-αSyn-108 and TAMRA-labelled NACore). Scale bar = 20 μm. (**C**) Tranfer free energy diluted phase-coacervate calculated from partition coefficients determined from microscopy experiments. (**D**) Gray value profiles of coacervate droplets from selected systems.

To determine whether α-synuclein and its variants partition into coacervate droplets or remain excluded, Alexa Fluor-647-labelled variants of α-synuclein were added to the coacervate emulsions and placed in a chambered glass slide to visualise them with confocal microscopy. Interestingly, distinct partitioning could be observed for different combinations of labelled protein and coacervates (fig. 2B and 2C). FL-αSyn accumulated at the interface of the coacervate droplets and the solution phase, which was particularly visible for pLys/pGlu and pLys/ATP systems. For all coacervates the average fluorescence inside the droplets (excluding the interface) was higher than in the surrounding dilute phase. The truncated variant αSyn-108 remained excluded from all coacervate droplets, and particularly for pLys/pGlu system for which the ratio of concentration inside/outside was lowest. Finally, the NACore fragment partitioned into RP3/polyU droplets, and very weakly into pLys/ATP droplets, but remained excluded from coacervates formed by pLys/pGlu.

The tendency of FL-αSyn to localise to the interface of coacervate droplets may stem from the fact that its disordered chain includes both charged/hydrophilic and hydrophobic regions. Both large, negatively charged RNAs and small, hydrophobic dyes, have been found to partition into pLys/ATP coacervates (44, 45). However, in an amphiphilic molecule, such as FL-αSyn, not all regions are preferentially taken up by the coacervate environment, resulting in a strong localization at the interface. Previous studies on partially unfolded proteins have also shown similar interfacial localization (55). Interfacial localization seems to be strongest for coacervates with relatively low molecular weight anionic components, such as ATP in pLys/ATP and pGlu in pLys/pGlu. Displacing these small anions with FL-αSyn in the coacervates results in a larger gain in entropy than displacing the large polyU in RP3/polyU coacervates. Some uptake of FL-αSyn inside the coacervates is possible for all coacervates tested (Figure 2C), and can be explained by an overall favourable interaction between FL-αSyn and one of the components in the coacervates (55–58). The negatively charged C-terminal domain appears crucial for both the uptake and the interfacial localization: the truncated αSyn-108 was systematically excluded from the droplets. The NAC core does not show any interfacial localization, presumably because it only contains the relatively hydrophobic core region of the protein. Instead, it is either sequestered and distributed homogeneously inside the coacervates or excluded, reflecting the favourable interactions between NACore and RP3, and the unfavourable interactions with pGlu most likely.

### Aggregation kinetics in the presence of coacervates

Upon phase separation most coacervate forming material (peptides and RNA in our case) is condensed into droplets, which are in equilibrium with the surrounding diluted phase (supernatant). The supernatant usually contains very low, but not negligible concentrations of the coacervate components. We used thioflavin T (ThT) assay to study the aggregation kinetics of the αSyn variants. To distinguish the influence of coacervate droplets from the soluble components in the supernatant, we performed experiments in the presence of droplets and control experiments with only the supernatant (separated from coacervate droplets by centrifugation) for each coacervate system (example for RP3/polyU is shown in fig. 3A, full traces shown in supplementary fig. S1). In addition, a reference experiment was performed without any droplets or soluble coacervate components.

**Fig. 3.**
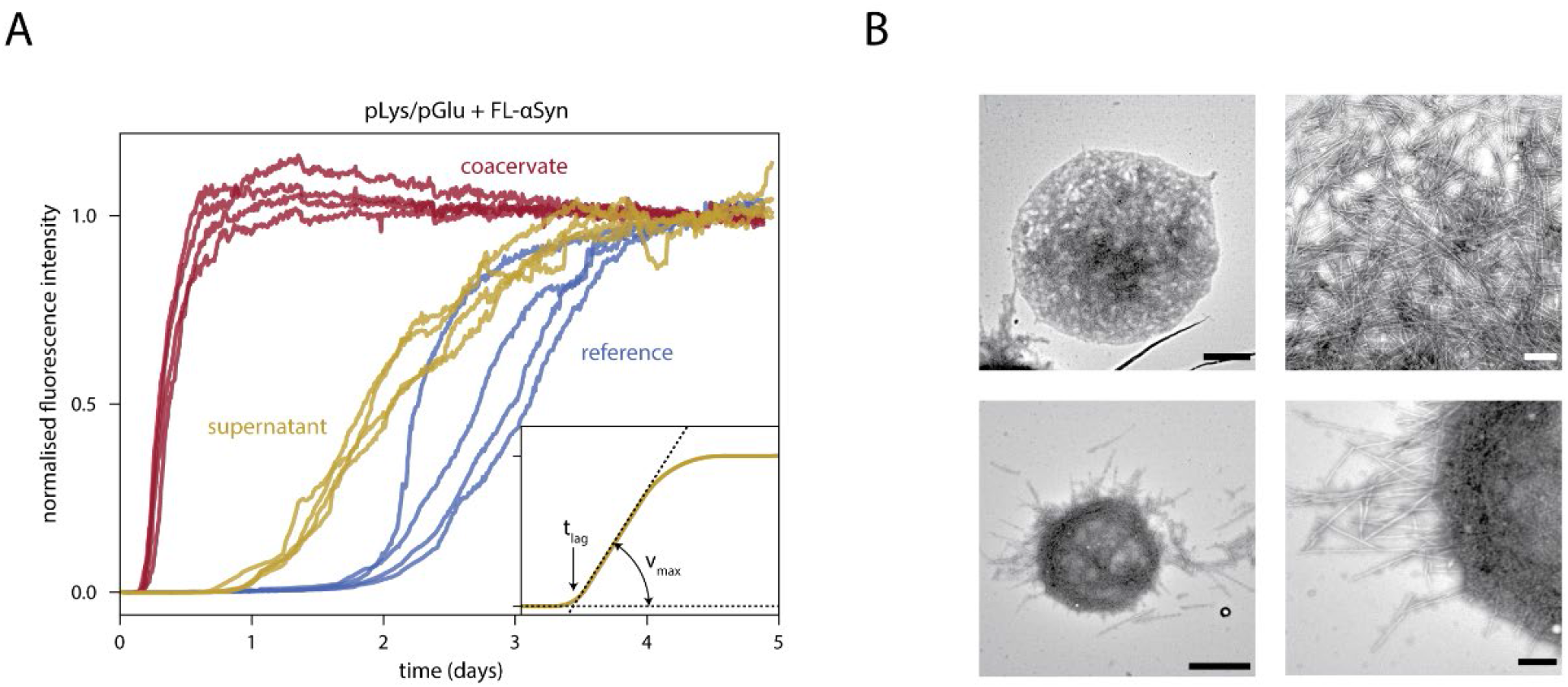
FL-αSyn aggregation in the presence of coacervates. (**A**) Aggregation traces for FL-αSyn: without coacervates (reference), with RP3/polyU supernatant and with RP3/polyU coacervates. (**B**) TEM images of aggregates formed in the presence of RP3/polyU coacervates. Scale bar: 1 μm for the images on the left side and 200 nm for the images on the right side.

Figure 3A shows a typical set of aggregation traces for one coacervate system and for one αSyn variant (FL-αSyn). It includes data for the reference sample without any coacervate material (blue traces), for the supernatant sample with diluted components shows faster aggregation (yellow traces) and for the sample with coacervate droplets (red traces). Before analysing the kinetics of aggregation further, we used TEM to determine whether the amyloid fibrils formed in the presence of droplets appear similar to the fibrils formed in solution. Figure 3B shows TEM images of FL-αSyn fibrils present inside and at the interface of coacervates droplets. The fibrils at the interface appear to be aligned, and some fibrils protrude into the surrounding solution. These results indicate that the fibrils can nucleate and grow in coacervate droplets. Moreover, analysis of the fibrils shown in fig. 3B revealed that there is no significant difference in thickness between these and fibrils formed in solution (supplementary fig. S2). Finally, we also purified the incubated samples of αSyn with coacervates by dissolving the coacervates at high salt concentration, depositing the fibrils on a TEM grid and rinsing the grid with Milli-Q water. All combinations of αSyn variants and coacervates show the same fibril appearance as in samples without coacervates (fig. 3 and supplementary fig. S2B).

To elucidate the role of condensates on the kinetics of aggregation, we plotted the distributions of both the lag times (tlag, which is predominantly determined by the primary nucleation rate), and the maximum slopes of the ThT-based aggregation curves (vmax, which is mostly determined by the elongation and secondary nucleation rate). As can be seen in fig. 4, the presence of each of the coacervates resulted in faster aggregation for FL-αSyn, even though the localization of FL-αSyn in these coacervates is not identical: in the case of RP3/polyU we observed a homogeneous distribution inside the droplets, while in the other two cases we observed a strong interfacial adsorption (fig. 1B).

The presence of RP3/polyU droplets mostly affect the lag phase of aSyn aggregation. With these droplets the lag phase was four times shorter than in controls with only supernatant and ten times faster than in reference solution, indicating that the amyloid nucleation rate was enhanced by the droplets. On the other hand, the maximum aSy aggregation rate in the presence of RP3/polyU droplets is comparable to the control samples with the RP3/polyU supernatant and to the reference sample without any coacervate material. Since these droplets concentrate FL-αSyn (fig. 2), it was expected that the growth rate inside the droplets is enhanced, as we discuss below.

**Fig. 4.**
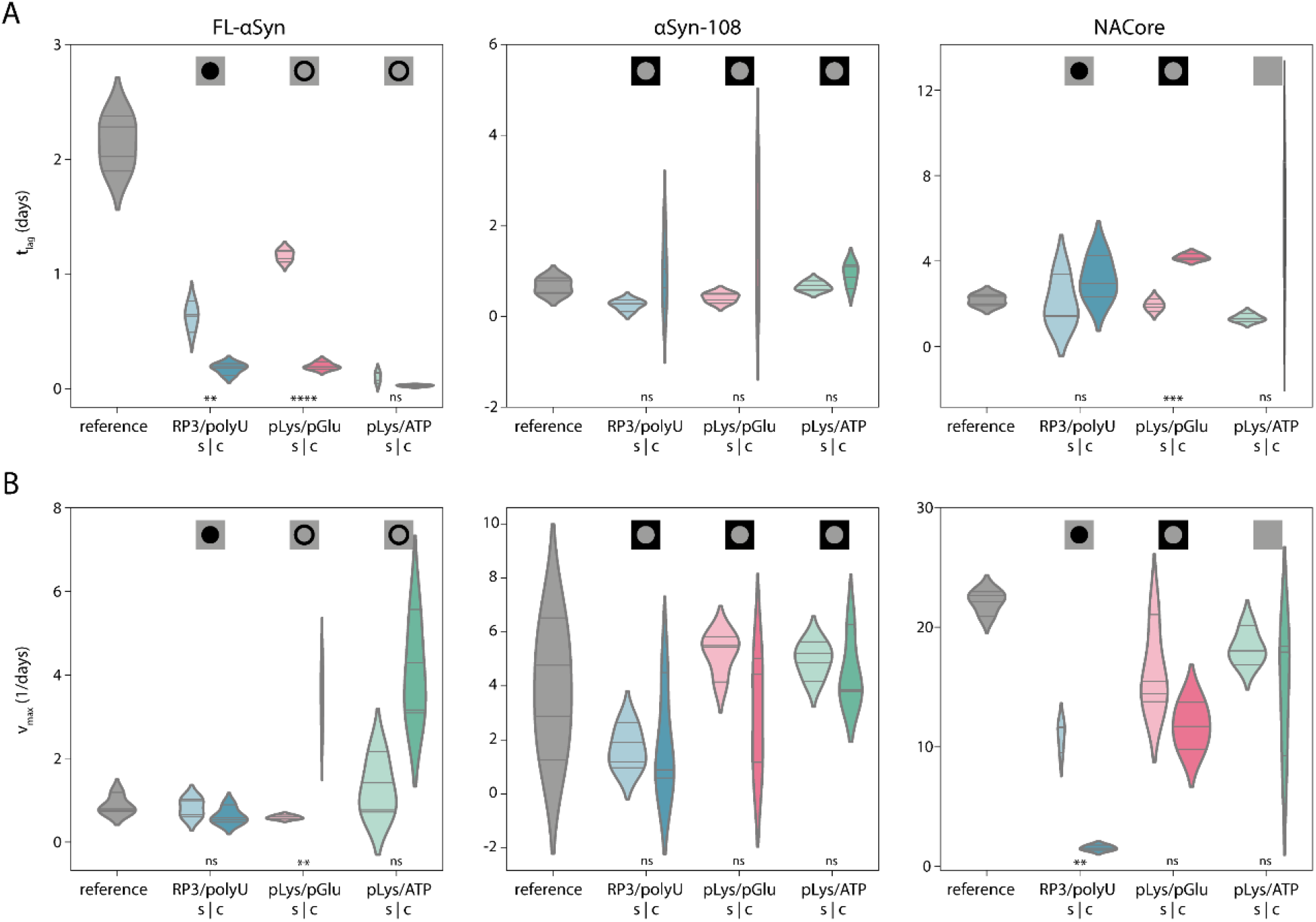
Analysis of aggregation kinetics. (**A**) Distribution of the lag times (*t*_lag_) for all protein variants and all coacervate systems (supernatant – s, coacervate – c) and for the reference sample. Symbol at the top indicate localisation of the corresponding variant as determined using fluorescence microscopy. Differences between samples were tested for statistical significance in coacervate droplets-supernatant control pairs. “ns” indicates values above 0.05, single asterisk indicates α<0.05, double asterisk - α<0.01, triple - α<0.001 and quadruple - α<0.0001. Violins plots were prepared using Gaussian kernels with bandwidth determined automatically using Scott’s method; density plots were cut at two bandwidth units past the extreme data points; violins are scaled to have the same area in supernatant-coacervate pairs. (**B**) Distribution of the maximum aggregation rates (*v*_max_) for all protein variants and all coacervate systems (colours and symbols as in A).

The fact that we observe a comparable growth rate despite a higher local concentration indicates that FL-αSyn is less aggregation-prone inside RP3/polyU coacervates. Possibly, one of the coacervate components can bind weakly to FL-αSyn monomers, oligomers or short fibrils and slow down amyloid growth. Nevertheless, the presence of droplets accelerated aggregation overall, in the sense that the time to complete aggregation was reduced, due to a shorter lag phase.

Different behaviour was observed for FL-αSyn aggregating in the presence of pLys/pGlu and pLys/ATP coacervates. The FL-αSyn in both the supernatant and the coacervate sample showed a faster onset of aggregation. Unlike in the case of RP3/polyU, the growth phase in the presence of pLys/pGlu and pLys/ATP droplets was significantly faster than in the reference sample, and also faster than in the presence of supernatant. This may be linked to the fact that pLys/pGlu and pLys/ATP systems seem to have a tendency to accumulate monomeric FL-αSyn at the interface of the droplets, thereby providing an alternative aggregation pathway, as we will discuss below. RP3/polyU system mostly accumulates FL-αSyn monomers inside and increase in concentration and altered environment affects mostly the primary nucleation rate.

The same experiments were performed for the αSyn-108 variant and the NACore peptide. Interestingly, the aggregation behaviour of the shorter variants was fundamentally different from the FL-αSyn. While the samples incubated in supernatant aggregated at comparable rate to the references for both αSyn-108 and NACore, the presence of droplets resulted in slower aggregation. Large spread of the aggregation parameters for αSyn-108 made it difficult to assess the significance of the effect, but for NACore it was clear that the peptide in samples with coacervates aggregated significantly slower than peptide in both the supernatant and reference samples. Presence of all types of droplets resulted in lag times that were longer than in the corresponding supernatant sample, although the spread was typically very large, which made only the lag time of pLys/pGlu droplets statistically significant. The presence of RP3/polyU droplets also resulted in significantly slower maximum aggregation rates than in supernatant, suggesting that the NACore peptides sequestered inside these droplets are less aggregation prone, for reasons we discuss below.

### Spatiotemporal mapping of the aggregation process

To find out if the faster and slower aggregation is related to the location where aggregation takes place, as the partitioning data (fig. 2B) seem to suggest, and to follow the spatiotemporal distribution of aggregates in the presence of coacervates, we developed an intramolecular FL-αSyn FRET probe, similar to the probe used before to study conformations of αSyn at a single molecule level (59, 60). The probe includes two fluorescent dyes close to the region responsible for β-sheet formation: Alexa Fluor 546 and Alexa Fluor 647 (fig. 5A). Upon aggregation, the dyes become fixed close to each other, which results in enhanced energy transfer (fig. S3). Experiments were performed in a similar fashion to the partitioning experiments, but the images were collected for several days and the samples were incubated at 37 °C. Collected fluorescence intensity images were used to create FRET efficiency maps, by calculating FRET efficiency for each pixel separately (fig. 5B).

**Fig. 5.**
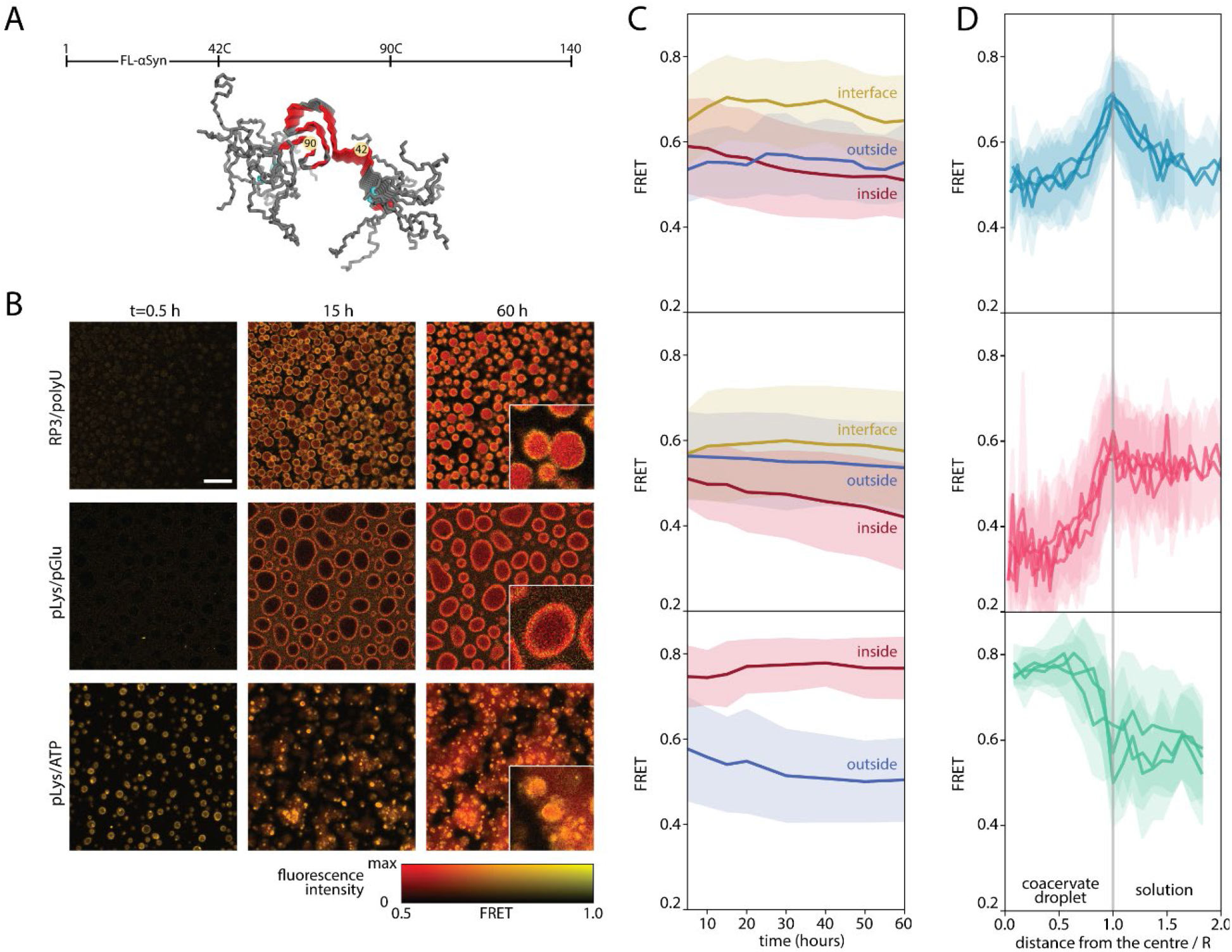
Aggregation monitored by FRET. (**A**) Positions of FRET labels in the FL-αSyn chain (PDB ID: 2N0A). (**B**) FRET maps of coacervate samples incubated with FRET-labelled FL-αSyn. Scale bar: 20 μm. Insets show 3 times enlarged part of the image at 60 h; experiment was performed in the presence of the FRET probe and 10 μM (RP3/polyU and pLys/ATP) or 40 μM (pLys/pGlu) concentration of non-labelled FL-αSyn. (**C**) Changes in FRET intensity in different areas of the coacervate systems over time. (**D**) FRET intensity radial profiles for coacervate droplets after 60 hours of incubation (distance from the droplet centre normalised by the diameter).

We observed distinct behaviour for the different coacervates. In the case of RP3/polyU the FRET signal increased throughout the entire coacervate droplet directly from the beginning, and was slightly enhanced at the interface, while it remained constant and low outside the droplets (fig. 5C and 5D). After 60 hours of incubation FRET signal increased inside the droplets, which suggests that the aggregates are formed inside the droplets, or at the interface and then move to the interior (fig. 5B).

For pLys/pGlu, the FRET efficiency was higher at the interface than in the surrounding solution, and interestingly, also higher than inside the droplets, even after 60 h. This suggests a more compact conformation and high concentration of FL-αSyn at the interface, potentially promoting faster nucleation into fibrils, which form at the surface of the droplets, but do not move towards the interior. In the case of pLys/ATP, aggregation starts immediately, with practically no lag phase (as also observed in the kinetic experiments, fig. 4). The highest FRET efficiency was observed inside the coacervate droplets, and the droplets seem to maturate over time, losing their liquid properties. Therefore, for the pLys/ATP system we also expect that the aggregates nucleate within the coacervate droplets, despite the enhanced concentration of monomeric αSyn at the interface, as observed in the partitioning experiments (fig. 2B). In FRET experiments, no accumulation of aggregated FL-αSyn could be observed at the interface of pLys/ATP coacervates. Presumably, even if aggregates nucleate at the interface, they immediately move to the interior of the droplets.

Similar observations were made using unlabelled FL-αSyn and ThT as a dye to stain the fibrils under the microscope (supplementary fig. S4A). In this case, presence of protein aggregates is simply indicated by high ThT fluorescence intensity. Direct analysis of the fluorescence intensities is complicated in this case, because free ThT also partitions into coacervates (supplementary fig. S4B). Nevertheless, we could observe significant increases in fluorescence intensity upon aggregation. In the presence of RP3/polyU and pLys/ATP droplets, aggregates were formed within the droplets resulting in irregular solid-like particles. In the presence of pLys/pGlu coacervates, the highest fluorescence intensity was observed for the coacervate interface, suggesting that aggregation is promoted by droplet interface, as observed also for the FRET probe.

### A kinetic model of protein aggregation accelerated or suppressed by condensates

Our microscopy experiments suggest that the presence of the coacervate droplets can affect α-synuclein aggregation process either through partitioning of the protein into coacervate droplets or through α-synuclein/coacervate interface interactions. To prove that these interactions can be also the reason of differences observed in the kinetics of the aggregation process, we developed and fitted kinetic models to our experimental data (fig. 5). Two separate models were developed to match the observations in fig. 2: (i) α-synuclein is excluded from or sequestered by the droplets and aggregation can take place both inside and outside the coacervate droplets (fig. 5A), (ii) α-synuclein is localized at the coacervate interface and heterogeneous nucleation followed by further aggregation can take place at the interface (fig. 5B). A detailed description of the differential equations for both models can be found in the supplementary information.

In the first case of exclusion or sequestration, we assumed that α-Syn monomers are freely exchanged between the dilute phase (which we call supernatant hereafter) that surrounds the coacervate droplets, and that the exchange of α-Syn between the supernatant and the condensed phase is much faster than the aggregation process itself. Consequently, local α-Syn concentrations are always equilibrated (i.e., the partition coefficient, as determined in fig. 2C, is constant). Aggregation of α-Syn can occur both in the supernatant and in the coacervate droplets and the rate constants of each step of the aggregation process (fig. 1C) can be different in both phases, which makes our approach different from previous aggregation models. Kinetic rate constants for the data obtained for α-Syn in supernatant were determined by fitting a simple aggregation model (for a one-phase system) and were further used as input for the supernatant phase when fitting parameters for the coacervate-containing samples (fig. 5C), thereby reducing the number of fit parameters. Finally, the fibrils are assumed to be immobile, in accordance with previous models (32): once formed they remain in the diluted or condensed phase.

Partitioning into the coacervate phase can accelerate aggregation because of increased local α-synuclein concentrations, but different rate constants for the aggregation process inside the coacervate droplets might mask this effect or further enhance it. In our experiments, we know the local concentration of α-synuclein from partitioning studies (fig. 2C), and can therefore deduce the additional influence of the coacervate environment on the rate constants. We note that the fits can only be used to obtain an order-of-magnitude estimate of the rate constants, as it is difficult to determine the separate contributions of each step in the aggregation process from a fit without comparing data collected for different α-synuclein concentrations. Nevertheless, we found significant differences between the rate constants in the supernatant and the coacervate phase. The fitted aggregation curves for RP3/polyU suggest that the primary nucleation rate of FL-αSyn inside the droplets is higher than in the supernatant, while the elongation and secondary nucleation rates are slightly lower, implying that the RP3/polyU coacervate environment has a destabilizing effect on monomeric FL-αSyn, but a stabilizing effect on oligomers and fibrils, under the assumption that all steps in the aggregation process are not diffusion limited. For NACore, which was also sequestered in RP3/polyU coacervates, we found a similar trend, although the model is not able to capture the very sharp onset of aggregation in some cases (supplementary fig. S5B).

The truncated αSyn-108 was excluded from all types of coacervates and, hence, we did not observe any significant difference in aggregation kinetics. Even in the presence of droplets, the aggregation process takes place predominantly in the supernatant, as there is hardly any αSyn-108 present inside the droplets. Therefore, we could not reliably determine the rate constants of aggregation for αSyn-108 in the coacervates.

The same is true for NACore in the presence of pLys/pGlu coacervates and pLys/ATP coacervates where we observed an overall suppressed aggregation, manifested in a longer lag time, but the monomers did not show significant sequestration. On the contrary, they are excluded from pLys/pGlu coacervates and indifferent to pLys/ATP coacervates. Therefore, a simple three-step model as shown in fig. 5A cannot explain the suppressed aggregation (fig. S5).

In other cases, we observed strong accumulation of the αSyn variants at the interface of the coacervates, rather than inside, which suggests a different mechanism of aggregation. Therefore, we developed a kinetic model to describe binding of the protein to the interface and further heterogeneous nucleation. The simplest model involving only binding and heterogeneous (primary) nucleation was not able to capture the very rapid aggregation observed in some cases. When we also allowed elongation and secondary nucleation to occur at the interface, we could capture the rapid global aggregation (fig. 5B). For pLys/ATP coacervates the primary nucleation rate constant at the interface is several orders of magnitude higher than in the supernatant, while other rate constants appear the same.. This can explain the very rapid onset of aggregation with virtually no lag time for pLys/ATP. For pLys/pGlu coacervates, primary nucleation and elongation seem unchanged, but the secondary nucleation rate constant is significantly faster at the interface, which explains the very rapid increase in ThT fluorescence after a lag phase (fig. 5B).

To confirm that the coacervate interface is crucial in enhancing the aggregation kinetics we have performed additional experiments in which we changed the amount of available surface area. In the first experiment we varied the amount of droplets-forming material. In the second experiment we centrifuged the coacervate dispersions before adding αSyn, causing the droplets to fuse (and thus reducing the available surface area). In both experiments we could observe that reducing the droplet surface area resulted in slower aggregation (supplementary fig. S6).

## Discussion

Our results show that condensates that are composed of non-aggregating material themselves can influence the aggregation of amyloidogenic proteins, such as α-synuclein, significantly and in a wide variety of ways. For FL-αSyn we observed an increase in aggregation rate for all systems. As suggested before by Weber and co-workers (32), this influence can be at least partially caused by higher local concentration of the aggregating protein inside coacervates. However, partition coefficients determined for the studied systems do not seem to explain the difference in the aggregation kinetics, unless we assume different aggregation rate constants inside the coacervate droplets and in the surrounding solution. Such differences would not be unexpected, as the more crowded and hydrophobic coacervate environment (31, 61, 62), rich also in functional groups that can interact with α-synuclein, affect the protein conformation and its tendency to aggregate. Moreover, our partitioning data combined with FRET microscopy indicate a different mechanism of accelerated aggregation for RP3/polyU and pLys/ATP (mainly due to enhanced rate constants for primary nucleation and growth inside the coacervates) on the one hand, and pLys/pGlu on the other (mainly due to enhanced rate constants of secondary nucleation at the interface).

We have also observed for the first time that the coacervate interface can serve as a nucleation site for protein aggregation. Relatively high apparent kinetic rate constants determined for FL-αSyn at the pLys/pGlu droplets interface may suggest that the coacervate droplets do not only serve as simple heterogenous catalysis nucleation sites, but they may also provide a distinct environment for protein aggregation or allow for a different aggregation mechanism to occur. It is interesting to note that very recently different behaviour has been observed for FL-αSyn under conditions that promote phase separation of FL-αSyn itself (i.e., in the presence of PEG and at high concentrations). Under such conditions, FL-αSyn forms liquid droplets that undergo maturation (a transition into solid aggregates) and this transition was found to be initiated at the centre of the droplets, suggesting that FL-αSyn droplets also provide a distinct environment in which the kinetic parameters of aggregation are altered, just like in the case of our pLys/pGlu droplets (63).

Furthermore, our results show that the influence of the coacervate droplets on aggregation kinetics depends on both the coacervate composition and the sequence/length of the aggregating protein. While aggregation of the full-length variant was accelerated in the presence of all coacervate systems, aggregation of the truncated variant, αSyn-108, was not significantly affected. This can be attributed to a different affinity of the full-length and the truncated αSyn variants to the coacervate material, and particularly to the positively charged components. The absence of the negatively charged C-terminal part in αSyn-108 makes this variant slightly positively charged at neutral pH (pI = 9.16), while FL-αSyn is strongly negatively charged (pI = 4.67). FL-αSyn has been shown before to aggregate faster in the presence of polycations in solution, and similar acceleration may occur inside coacervates or at their interface (64).

Another interesting observation is that pLys/pGlu and pLys/ATP affect the aggregation process differently, even though they both contain pLys. The reason for this difference is the binding strength of the counterions present in these coacervate droplets. ATP has fewer negative charges and binds less strongly to pLys than pGlu, which is evidenced by the lower critical salt concentration of pLys/ATP droplets. As a result, FL-αSyn can displace ATP more easily than pGlu and bind more strongly to pLys. We hypothesize that stronger binding of the negatively charged tail of αSyn makes the protein more prone to aggregation, similar to previous reports [REF]. In addition, the weaker interaction of ATP compared to pGlu leads to a lower viscosity inside the pLys/ATP condensates, which facilitates movement of aggregates and FL-αSyn bound to pLys inside the droplets.

Finally, coacervate droplets are also able to slow down aggregation, which was most prominent for NACore peptide. This may be explained by sequestration of free peptides and small oligomers inside the coacervate (fig. 2), in relatively stable conformation, not prone to rapid aggregation. Surprisingly, in the case of pLys/pGlu droplets, where labelled NACore peptide remained excluded from the droplets, the aggregation was also slowed down. It is possible that while free peptides were excluded, small oligomers, which form in early stages of the aggregation process, are sequestered by the droplets and prevented from further growth. However, proving this is impracticable, because any action to separate the droplets from the supernatant will most likely disrupt such oligomers.

In conclusion, we show that pre-existing liquid condensates can affect protein amyloid formation in vitro, both accelerating and slowing down the reactions. We expect that the same process can happen in living cells, which contain multiple membraneless organelles, formed upon LLPS. By sequestering amyloidogenic proteins, such biological condensates may prevent protein aggregation, but it is also possible that they can function as heterogenous nucleation sites. This provides a new perspective on the early stages of amyloid formation by α-synuclein (and protein aggregation in general) in the complex cellular environment.

## Materials and Methods

### Reagents

Poly-L-lysine hydrobromide (MW 15-30 kDa), adenosine 5’-triphosphate (ATP) disodium salt, polyuridylic acid potassium salt, buffers and thioflavin T (ThT) were purchased from Sigma-Aldrich. RP3 (RRASLRRASLRRASL-NH2), and NACore (GAVVTGVTAVA) peptides were purchased from CASLO ApS (Denmark). Labelled NACore peptide was synthesised on solid phase using standard Fmoc peptide synthesis strategy. Poly-D,L-lysine hydrobromide (MW = ca. 21 kDa) and poly-D,L-glutamic acid sodium salt (MW = ca. 15 kDa) were purchased from Alamanda Polymers (USA). Alexa Fluor maleimides were purchased from Thermo Fisher. Poly-L-lysine grafted with poly(ethylene glycol) (PLL-g-PEG) was purchased from SuSoS AG (Switzerland). All aqueous solutions were filtered before use using Acrodisc 0.2 μm nylon syringe filters (Sigma-Aldrich) or Pierce cellulose acetate filter spin cups with 0.45 μm pore size (Thermo Fisher).

### Protein preparation and labelling

Wild-type FL-αSyn, αSyn-108 and the cysteine mutants were expressed and purified as previously described (65). Purified proteins were stored at a concentration of ~200 μM in 10 mM Tris-HCl, pH 7.4 at −80 °C, supplemented with 1 mM DTT for the cysteine mutants. Single labelled proteins were labelled according to the dye manufacturer procedures. For labelling of double-cysteine mutant (42C 90C), the first labelling step (with donor dye) was performed according to the dye manufacturer procedures, using 1:1 protein to dye ratio. Subsequently, the protein was incubated with pre-washed Activated Thiol–Sepharose 4B (Cytiva, USA) for 1 hour, rotating in the dark at 4 °C. Next, the resin was washed with several volumes of 10 mM Tris-HCl, pH 7.4, followed by elution of single and double labelled αSyn using buffer containing 25 mM DTT. Eluted fractions were pooled, concentrated to about 0.5 mL and desalted. Triple excess of acceptor dye was added to the concentrated protein and the solution was incubated for 1 hour at room temperature. Unbound dye was removed using Amicon Ultra-4/15 centrifugal filters with suitable MWCO.

Protein solutions were filtered using Pierce cellulose acetate filter spin cups (Thermo Fisher) before every aggregation kinetics assay and concentration was determined based on absorbance (ϵ = 5600 M^-1^ cm^-1^ for wild-type αSyn and ϵ = 1400 M^-1^ cm^-1^ for αSyn(1–108)).

### Preparation of modified glass slides

All glass slides used for microscopy were modified according to the following procedure. First, the slide was washed thoroughly with milliQ water. Subsequently, the surface intended to be modified was cleaned with oxygen plasma and 0.01 mg/ml solution of PLL-g-PEG in 10 mM HEPES buffer (pH 7.4) was applied on the glass immediately after plasma treatment. Glass was incubated with the PLL-g-PEG solution for 2 hours at room temperature. Subsequently it was rinsed 3 times with 10 mM HEPES buffer (pH 7.4), 3 times with milliQ water and dried with pressurised air. Modified slides were stored at room temperature and used within one week.

### Partitioning of labelled protein

Localisation of labelled proteins was studied using confocal microscopy. Leica SP8x confocal microscope equipped with 40x magnification water-immersion objective was used. Samples were placed in 18-well chambered glass coverslips (Ibidi GmbH, Germany), previously modified with PLL-g-PEG. Partition coefficients were determined by calculating ratio of fluorescence intensity in the condensed phase to fluorescence intensity in the outer phase (average intensity values from at least 10 droplets and from outer phase of similar area was used). Background signal of coacervate sample without labelled protein was subtracted separately for condensed and supernatant.

### ThT aggregation kinetics assays

To estimate the aggregation kinetic parameters, we have performed standard Thioflavin T (ThT) aggregation assays. Upon binding to β-sheets, ThT fluorescence intensity increases by several orders of magnitude and the changes of fluorescence in the solutions of aggregating protein are proportional to the amount of aggregate formed ([M]).

Aggregation assays were performed under following conditions: 50 mM HEPES, 100 mM NaCl, 100 μM EDTA, 20 μM ThT, and 40 μM FL-αSyn or αSyn(1–108), or 160 μM of NACore. All aggregation assay were performed in non-binding 384-well plates with black walls (Greiner Bio-One GmbH, Austria), at 37 °C. To prevent evaporation, wells in the two outer rows were always filled with water and the plate was sealed with film. Measurements were performed using Tecan Spark or Tecan Infinite M200 microplate reader. Fluorescence intensity was recorded every 12 minutes using bottom readout with continuous linear shaking in between. Excitation and emission wavelength range was controlled using monochromators for Tecan Infinite M200 (respectively 440 nm with 9 nm bandwidth and 480 nm with 20 nm bandwidth) or filters for Tecan Spark (respectively 430 nm with 20 nm bandwidth and 460 nm with 20 nm bandwidth). To extract the basic kinetic parameters (tlag and vmax) from the ThT fluorescence traces we fitted simple aggregation model (as described in the supplementary information) and used the maximum slope of the curve as vmax and the intersection of line going through the max slope point and the baseline was used as tlag (see inset in fig. 3A).

### Preparation of samples and transmission electron microscopy

Samples after the ThT aggregation kinetics assay in 384-well plates were used for electron microscopy experiments. In order to dissolve the coacervate material and separate the α-Syn aggregates, sodium chloride solution was added to the selected wells, to final concentration of 300 mM sodium chloride. After incubation for 5 minutes at room temperature, plate was centrifuged for 10 minutes at 1000 rcf. Subsequently, solution was gently collected from the selected wells and 50 μl of milliQ water was added. The plate was centrifuged again with the same settings and again solution was gently collected from the selected wells. Any precipitate from the selected wells was resuspended in 20 μl of milliQ water, and subsequently 2 μl of the suspension was transferred onto a TEM grid (EM-Tec formvar carbon support film on copper, 300 square mesh, Micro to Nano, the Netherlands). Samples were blotted with filter paper, stained with 1.5 μl of 2% (w/w) sodium phosphotungstate solution (adjusted to pH 7.4), washed with 2 μl of water left to dry overnight. Imagining was performed using JEOL JEM-1400 FLASH.

### Intramolecular FRET experiments

FRET experiment was performed using Leica SP8x confocal microscope equipped with 40x magnification water-immersion objective. Samples were placed in 18-well chambered glass coverslips (Ibidi GmbH, Germany), previously modified with PLL-g-PEG and the whole setup was incubated at 37 °C during the experiment. FRET probe was added at 0.01 ratio to the non-labelled FL-αSyn (0.1 or 0.4 μM and 10 or 40 μM respectively), other components remained the same as for the ThT aggregation kinetics assay. Samples were excited at 488 nm and the emission was recorded at 515-530 nm for the donor and 590-610 nm for the acceptor. Fluorescence intensity images were saved in 8-bit 512×512 pixels format. FRET value was calculated for each pixel using the following formula:

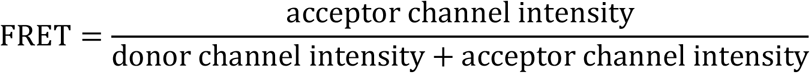

The FRET value was not determined if intensities for both channels were lower than 2, which is close to the detector dark count. A 512×512 array of FRET intensities was further converted into an 8-bit 512×512 image and visualized using custom hue/brightness 2D colour map.. Hue corresponds to the FRET value scaled from 0.5 to 1 (pixels with values below 0.5 have the same hue as pixel with FRET equal 0.5). Brightness is proportional to the sum of fluorescence intensity for both channels, scaled from 0 to the value for 95 percentile in the image collected after 60 h (pixels with higher intensity have the same “max” brightness).

The FRET experiment in bulk (fig. S3) was performed using 0.4 μM concentration of the FRET probe and 40 μM concentration of non-labelled FL-αSyn and incubated at 37 °C in a Eppendorf tube. Fluorescence spectra were measured using JASCO FP-8300ST spectrofluorometer.

### Statistical analysis

Microscopy images were analysed using FIJI distribution of ImageJ. Error bars and error ranges of transfer energies and FRET plots were determined using standard deviations of pixel intensity values within selected range. Plots in fig. 5C were prepared by manually selecting parts of the image. Plots in fig. 5D were prepared using radial profile angle plugin for ImageJ. Violin plots were prepared according to the description under fig. 4. Fitting of the aggregation kinetic models in fig. 6C was performed using basinhopping function from scipy.optimize library in python.

**Fig. 6.**
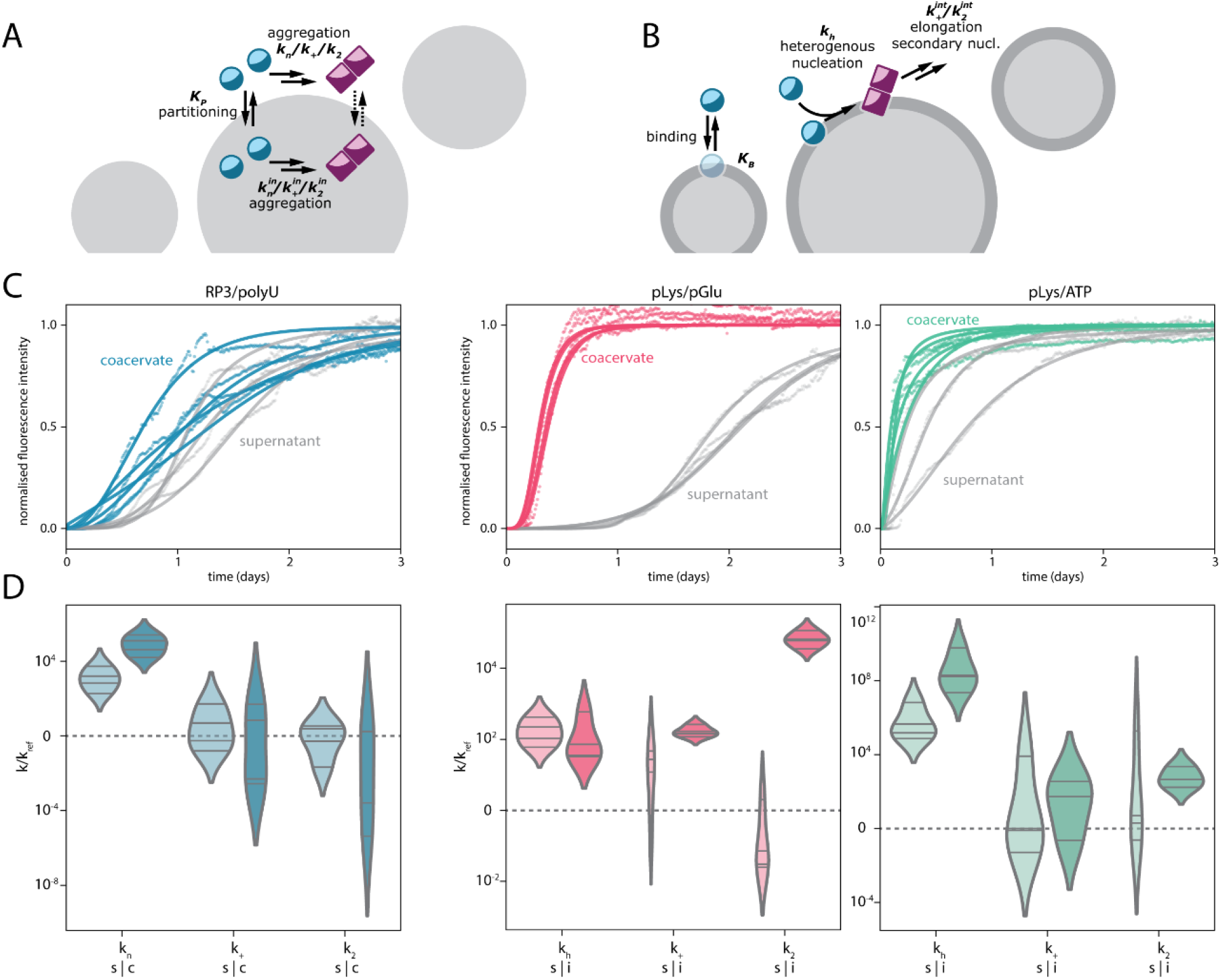
Fitting of the aggregation models. (**A**) Schematic depiction of the aggregation-in-droplets model. (**B**) Schematic depiction of the interface-aggregation model. (**C**) Fits to the experimental data for RP3/polyU with FL-αSyn for the aggregation-in-droplets model and fits to the experimental data for pLys/pGlu and pLys/ATP with FL-αSyn for the interface-aggregation model. (**D**) Resulting aggregation kinetic rates, for the diluted-supernatant phase (s) and for the coacervate/interface phase (c/i), normalised by values for reference sample (without coacervate components). Violin plots were prepared analogously to plots in fig. 4.

## Author contributions

T.L., W.T.S.H., E.S. generated ideas and wrote the manuscript. T.L. designed and carried out experiments. S. L. designed and wrote the model part with input from C. A. W. H.S. and L.S. helped with initial liposome preparation. All authors have discussed the results and approved the final version of the manuscript.

## Additional information

Supplementary information containing a detailed kinetic model, and supplementary figures is available.

## Acknowledgements

This work was supported financially by the Netherlands Organization for Scientific Research (NWO. The authors would like to thank K.A. van Leijenhorst-Groener (University of Twente) for purification of the αSyn variants and G.-J. Janssen (Radboud University) for help with TEM measurements. MAAF and MMAEC acknowledge the Dutch Parkinson’s disease foundation “Stichting ParkinsonFonds” for their support for the development of the FRET probe.

