## Supplementary information for "Biomolecular condensates can both accelerate and suppress aggregation of α-synuclein"

### Supplementary Text

#### Basic aggregation model

Typically for many amyloidogenic proteins, α-synuclein aggregation process may be considered an autocatalytic process. A simple yet accurate model of α-synuclein aggregation involves three basic reactions: (i) primary nucleation of fibres from α-synuclein monomers, (ii) elongation of fibres by attaching monomers to one of the fibre ends, (iii) secondary nucleation catalysed by fibres:

$$r_{\text{primary nucleation}} = k_{\text{n}} \cdot [S]^{n}$$

$$r_{\text{elongation}} = k_{+} \cdot [S] \cdot 2 \cdot [P]$$

$$r_{\text{secondary nucleation}} = k_{2} \cdot [S]^{n_2} \cdot [M]$$

where $k_{\text{n}}$, $k_{+}$, $k_2$ are the reaction rates of the corresponding reactions, $n$ and $n_2$ are the nucleation numbers of primary and secondary nucleation (the lowest number of oligomers required to form a fibre nucleus), and $[S]$, $[P]$ and $[M]$ are the concentration of monomers, concentration of fibres (so $2 \cdot [P]$ reflects the number concentration of fibril ends) and concentration of monomeric units incorporated in fibres (proportional to fibre mass concentration and the surface available for secondary nucleation catalysis).

From this a set of differential equations describing concentration changes in the system can be derived:

$$\frac{d[S]}{dt} = -n \cdot k_{\text{n}} \cdot [S]^{n} - k_{+} \cdot [S] \cdot 2 \cdot [P] - n_2 \cdot k_2 \cdot [S]^{n_2} \cdot [M]$$

$$\frac{d[P]}{dt} = k_{\text{n}} \cdot [S]^{n} + k_2 \cdot [S]^{n_2} \cdot [M]$$

$$\frac{d[M]}{dt} = n \cdot k_{\text{n}} \cdot [S]^{n} + k_{+} \cdot [S] \cdot 2 \cdot [P] + n_2 \cdot k_2 \cdot [S]^{n_2} \cdot [M]$$

Solving this set of equations provide a kinetic trace of the aggregation process. Fitting the solution to the experimentally measured concentration of one of the species provides information about the protein aggregation rates.

**Development and validation of protein aggregation model in coacervate systems**

In case of partitioning into the coacervate droplets, the concentrations of monomer in the diluted and in the condensed phase is determined by the partitioning constant:

$$K_{\mathrm{P}} = \frac{[S]_{\mathrm{cond}}}{[S]_{\mathrm{dil}}}$$

where $K_P$ is the partitioning constant and $[S]_{\mathrm{cond}}$ and $[S]_{\mathrm{dil}}$ are the concentrations of the monomer in the condensed and the diluted phase respectively. Taking into account the equation describing the mass balance of monomers in the system:

$$[S]_{\mathrm{tot}} = [S]_{\mathrm{dil}} \cdot \frac{R}{1+R} + [S]_{\mathrm{cond}} \cdot \frac{1}{1+R}$$

where $R$ is the ratio of diluted phase volume to the condensed phase volume, we can write equations describing the concentrations of the monomers in the diluted and in the condensed phase:

$$[S]_{\mathrm{dil}} = \frac{1+R}{R+K_{\mathrm{P}}} \cdot [S]_{\mathrm{tot}} = \xi \cdot [S]_{\mathrm{tot}}$$

$$[S]_{\mathrm{cond}} = K_{\mathrm{P}} \cdot \frac{1+R}{R+K_{\mathrm{P}}} \cdot [S]_{\mathrm{tot}} = K_{\mathrm{P}} \cdot \xi \cdot [S]_{\mathrm{tot}}$$

where $\xi = \frac{1+R}{R+K_{\mathrm{P}}}$, which further lead to a set of differential equations describing aggregation process in the coacervate system with monomer partitioning:

$$\frac{d[S]_{\mathrm{tot}}}{dt} = \left(\frac{R}{1+R}\right)\left[-nk_{\mathrm{n}}(\xi[S]_{\mathrm{tot}})^{n} - 2k_{+}\xi[S]_{\mathrm{tot}}[P]_{\mathrm{dil}} - n_2k_2(\xi[S]_{\mathrm{tot}})^{n_2}\,[M]_{\mathrm{dil}}\right] + \left(\frac{1}{1+R}\right)\left[-nk_{\mathrm{n_{cond}}}(K_{\mathrm{P}}\,\xi[S]_{\mathrm{tot}})^{n} - 2k_{+_{\mathrm{cond}}}K_{\mathrm{P}}\xi[S]_{\mathrm{tot}}[P]_{\mathrm{cond}} - n_2k_{2_{\mathrm{cond}}}(K_{\mathrm{P}}\xi[S]_{\mathrm{tot}})^{n_2}\,[M]_{\mathrm{cond}}\right]$$

$$\frac{d[P]_{\mathrm{dil}}}{dt} = k_{\mathrm{n}} \cdot (\xi[S]_{\mathrm{tot}})^{n} + k_2 \cdot (\xi[S]_{\mathrm{tot}})^{n_2}\,[M]_{\mathrm{dil}}$$

$$\frac{d[M]_{\mathrm{dil}}}{dt} = nk_{\mathrm{n}}(\xi[S]_{\mathrm{tot}})^{n} + 2\,k_{+}\xi[S]_{\mathrm{tot}}\,[P]_{\mathrm{dil}} + n_2k_2(\xi[S]_{\mathrm{tot}})^{n_2}\,[M]_{\mathrm{dil}}$$

$$\frac{d[P]_{\mathrm{cond}}}{dt} = k_{\mathrm{n_{cond}}}\,(K_{\mathrm{P}}\xi[S]_{\mathrm{tot}})^{n} + k_{2_{\mathrm{cond}}}(K_{\mathrm{P}}\,\xi[S]_{\mathrm{tot}})^{n_2}\,[M]_{\mathrm{cond}}$$

$$\frac{d[M]_{\mathrm{cond}}}{dt} = nk_{\mathrm{n_{cond}}}(K_{\mathrm{P}}\xi[S]_{\mathrm{tot}})^{n} + 2k_{+_{\mathrm{cond}}}K_{\mathrm{P}}\xi[S]_{\mathrm{tot}}[P]_{\mathrm{cond}} + n_2k_{2_{\mathrm{cond}}}(K_{\mathrm{P}}\xi[S]_{\mathrm{tot}})^{n_2}[M]_{\mathrm{cond}}$$

Again, similarly to the more simple case of aggregation in homogenous solution, solving the equations yields aggregation kinetic trace for both the diluted and the condensed phase.

Another model was developed for a case where aggregation-prone protein accumulates in the coacervate-diluted phase interface. Binding of the monomers to the coacervate interface can be described by equation:

$$K_{\mathrm{B}} = \frac{[S]_{\mathrm{int}}}{[S]_{\mathrm{dil}} \cdot [I]}$$

where $K_{\mathrm{B}}$ is the binding constant, $[S]_{\mathrm{int}}$ is the concentration of interface-bound monomers and $[I]$ is the concentration of available binding sites ($[I] = [I]_{\mathrm{tot}} - [S]_{\mathrm{int}}$). Again, taking into account the mass balance equation for monomers, we can write equations describing the concentration of free and surface-bound monomers. Since the aggregation reaction occurs now only in the diluted phase (or in the interface, which is treated as a part of the diluted phase), we can omit the change of volume:

$$[S]_{\mathrm{dil}} = \frac{-1 - K_B[I]_{\mathrm{tot}} + [S]_{\mathrm{tot}}K_{\mathrm{B}} + \sqrt{\left(1 + K_{\mathrm{B}}([I]_{\mathrm{tot}} - [S]_{\mathrm{tot}})\right)^2 + 4K_{\mathrm{B}}[S]_{\mathrm{tot}}}}{2K_{\mathrm{B}}}$$

$$[S]_{\mathrm{int}} = \frac{K_{\mathrm{B}}[I]_{\mathrm{tot}}[S]_{\mathrm{dil}}}{1 + K_{\mathrm{B}}[S]_{\mathrm{dil}}}$$

We assume that the surface can act as a nucleation site, requiring one monomer from the surface and one monomer from the solution to react. If we further assume that the fibres formed at the interface can grow by attaching monomers from the solution, they can participate in secondary nucleation and that they remain attached to the interface, we can write a set of differential equations for this system:

$$\frac{d[S]_{\mathrm{tot}}}{dt} = -nk_{\mathrm{n}}[S]_{\mathrm{dil}}{}^{n} - 2k_{+}[S]_{\mathrm{dil}}([P]_{\mathrm{dil}} + [P]_{\mathrm{int}}) - n_2k_2[S]_{\mathrm{dil}}{}^{n_2}([M]_{\mathrm{dil}} + [M]_{\mathrm{int}}) - 2k_{\mathrm{h}}[S]_{\mathrm{dil}}[S]_{\mathrm{int}}$$

$$\frac{d[P]_{\mathrm{dil}}}{dt} = k_{\mathrm{n}}[S]_{\mathrm{dil}}{}^{n} + k_2[S]_{\mathrm{dil}}{}^{n_2}[M]_{\mathrm{dil}} + n_2k_2[S]_{\mathrm{dil}}{}^{n_2}[M]_{\mathrm{int}}$$

$$\frac{d[M]_{\mathrm{dil}}}{dt} = nk_n[S]_{\mathrm{dil}}{}^{n} + 2k_+[S]_{\mathrm{dil}}[P]_{\mathrm{dil}} + n_2k_2[S]_{\mathrm{dil}}{}^{n_2}([M]_{\mathrm{dil}} + [M]_{\mathrm{int}})$$

$$\frac{d[P]_{\mathrm{int}}}{dt} = k_{\mathrm{h}}[S]_{\mathrm{dil}}[S]_{\mathrm{int}}$$

$$\frac{d[M]_{\mathrm{int}}}{dt} = 2k_{\mathrm{h}}[S]_{\mathrm{dil}}[S]_{\mathrm{int}} + 2k_+[S]_{\mathrm{dil}}[P]_{\mathrm{int}}$$

where $k_{\mathrm{h}}$ is the reation rate constant of the interface-catalysed nucleation and, for clarity, $[S]_{\mathrm{dil}}$ and $[S]_{\mathrm{int}}$ symbols were used instead of full equations dependent on $[S]_{\mathrm{tot}}$.

### Supplementary Figures

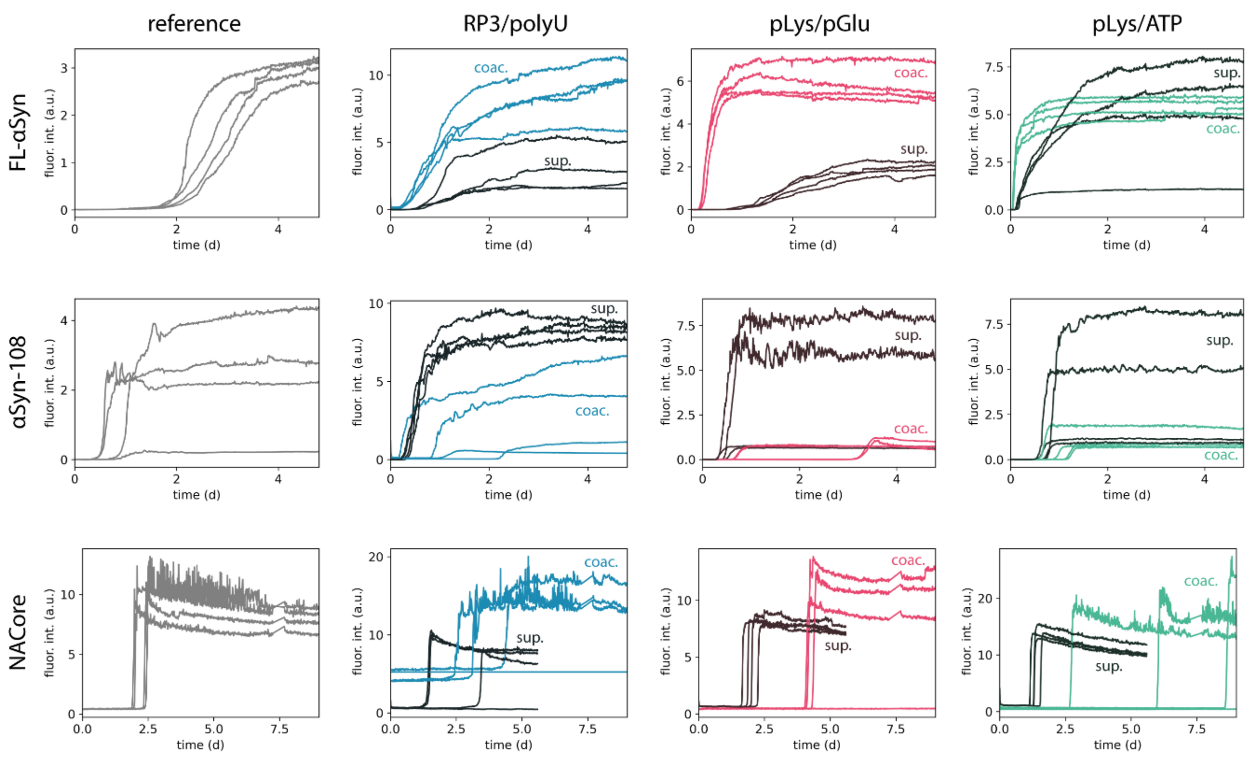


**Fig. S1.** Aggregation traces (ThT fluorescence intensity) of different αSyn variants in buffer (blank, grey traces), in the presence of coacervates (coloured traces), or in the presence of coacervate supernatants (dark traces).

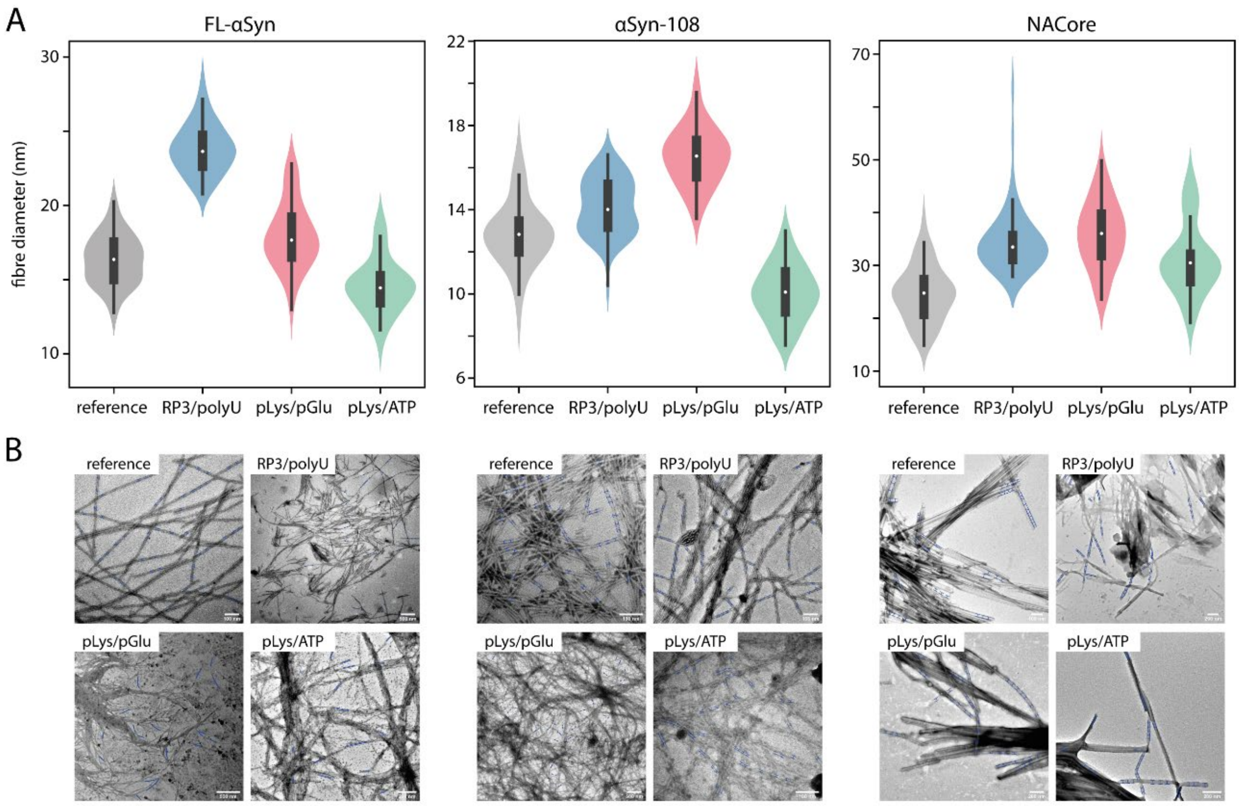


**Fig. S2.** (**A**) Distribution of fibre thickness formed by different αSyn variants in the absence (blank) or presence of coacervate systems (n=50). (**B**) TEM images of the fibres formed by different αSyn variants in the absence (blank) or presence of coacervate systems. Blue marks indicate places where diameter was measured.

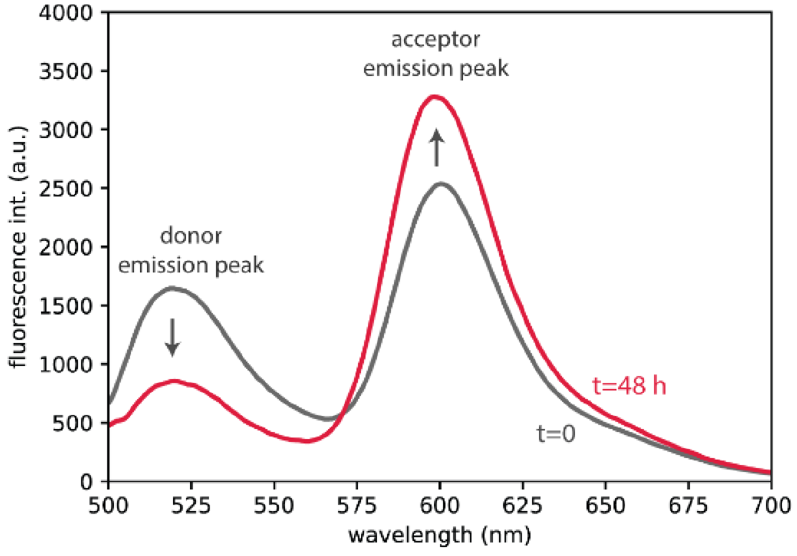


**Fig. S3.** Fluorescence spectra of the FL-αSyn-based FRET probe in solution (in bulk), shortly after preparing the solution (t=0) and after 48 hours of incubation at 37 °C (t=48 h).

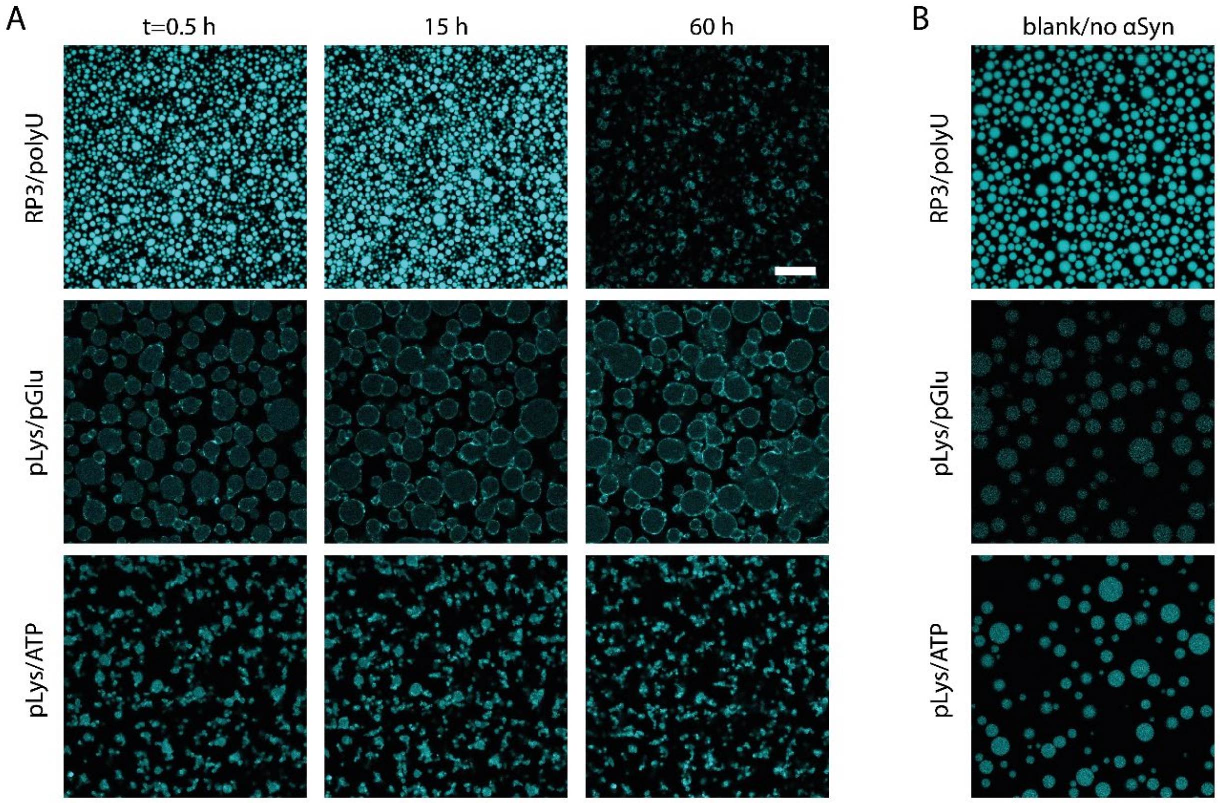


**Fig. S4.** (**A**) ThT aggregation assay under confocal microscope of FL-αSyn in presence of different coacervate systems. (**B**) Partitioning of ThT into coacervate systems (without added FL-αSyn).

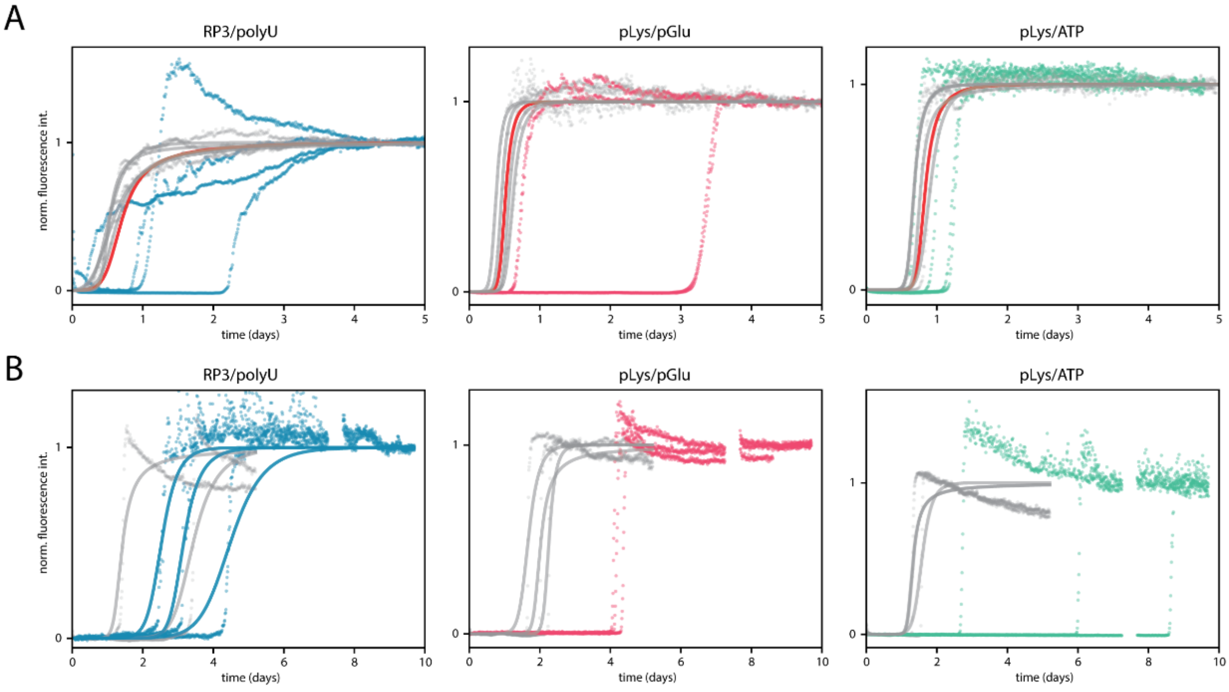


**Fig. S5.** (**A**) Aggregation of αSyn-108 in the presence of different systems; supernatant traces with fitted curves are shown in grey (and average in red) and coacervate traces are shown in colour. (**B**) Aggregation of NACore in the presence of different systems; supernatant traces with fitted curves are shown in grey and coacervate traces are shown in colour. Proposed models for aggregation in the presence of coacervate systems can explain similar aggregation kinetics in the presence of droplets without partitioning, but fails to explain slower aggregation in the presence of droplets with low to moderate partitioning.

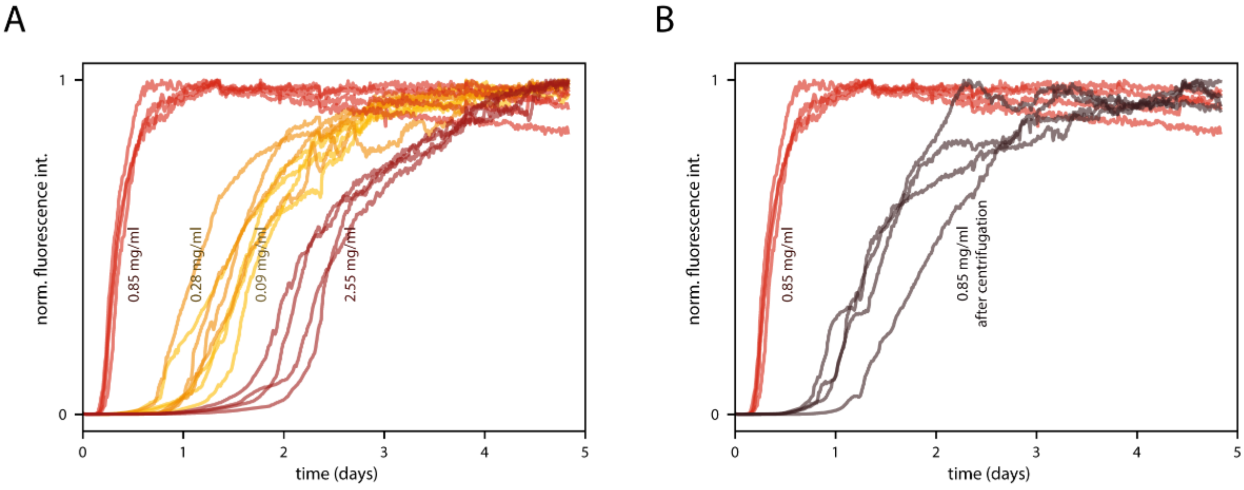


**Fig. S6.** (**A**) Aggregation traces of FL-αSyn in the presence of different amount of pLys/pGlu coacervates. (**B**) Aggregation traces of FL-αSyn in the presence of coacervates dispersed in solution and fused at the bottom of the plate after centrifugation.

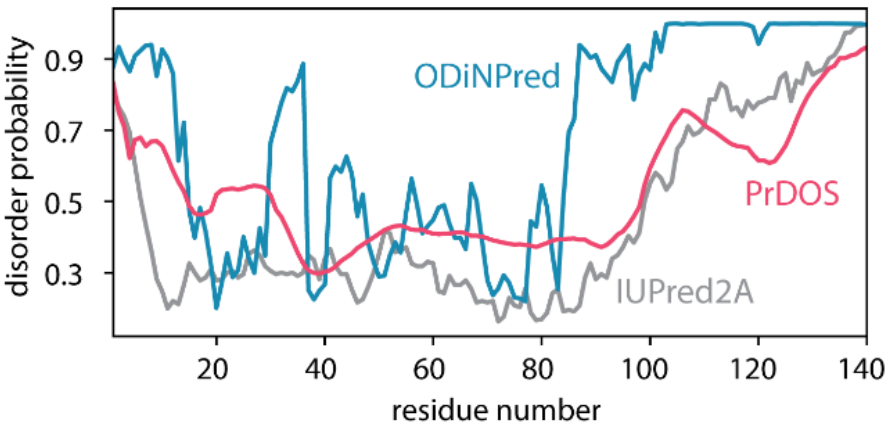


**Fig. S7.** Comparison of predicted disorder probability for FL-αSyn using different predictors: ODiNPred (*1*), PrDOS (*2*) and IUPred2A (*3*).
